# Low-Rank Tensor Decomposition for Cross-Bispectral Analysis of EEG Data

**DOI:** 10.64898/2025.12.21.695795

**Authors:** Dionysia Kaziki, Andreas K. Engel, Guido Nolte

## Abstract

Cross-bispectral measures provide a rich description of interactions in EEG signals, but their third-order tensor structure poses substantial challenges for interpretation and dimensionality reduction. We introduce a low-rank tensor decomposition framework specifically designed for cross-bispectral EEG data. The model expresses the bispectrum as a product of a single spatial mixing matrix and a compact source-interaction tensor, yielding a structured and interpretable representation of the data. Unlike classical tensor decompositions such as Tucker and its special case PARAFAC, where either the core tensor is unconstrained or restricted to rank-one terms, our formulation incorporates a source-space interpretation directly through the tensor *Q*_*mnp*_, making the decomposition well suited for identifying dominant coupling components. After obtaining the low-rank representation, spatial demixing is performed using MOCA to derive clear and interpretable source maps. Through simulations, we show that the method consistently retrieves the dominant structure of the bispectrum across a wide range of conditions, including changes in SNR, dipole orientation, amplitude balance, and source complexity. Applied to resting-state EEG, the method produces anatomically plausible parietal generators associated with the strongest alpha-band bispectral interactions. Overall, the framework provides a principled and computationally efficient approach for reducing and interpreting cross-bispectral EEG data, offering source-level insights that are difficult to obtain using standard tensor factorization techniques.

## 1 Introduction

In recent years, brain research has increasingly recognized the complexity of neural activity. Traditional methods for analyzing brain data, such as electroencephalography (EEG) and magnetoencephalography (MEG) recordings, primarily focus on frequency and time domains, often overlooking the spatial dimension. A more sophisticated approach involves structuring neural signals as multi-way arrays indexed by spatial, frequency, and temporal components, providing a fuller, more nuanced picture of brain activity [1]. This approach is essential for exploring interactions between different components of brain activity and advancing our understanding of neural dynamics. EEG/MEG oscillatory activity is complex and cannot simply be considered as noise. Instead, it involves non-linear interactions that require sophisticated analytical approaches [2]. Conventional methods, such as spectral analysis, often miss important aspects of these interactions [3, 4].

Cross-frequency coupling, the interaction between neural oscillations at different frequencies, facilitates coordination between neural regions operating at different temporal scales. Cross-frequency coupling has emerged as a fundamental mechanism of brain coordination, facilitating the integration of information across spatial and temporal scales and playing a crucial role in neural communication [5]. Furthermore, novel methods that capture cross-frequency synchronization, have been developed to address gaps left by traditional techniques that focus only on synchronization within the same frequency band [6, 7, 8, 9, 10]. Advanced techniques like higher-order spectral measures, such as bispectrum and bicoherence, are better suited to quantify stable phase relationships and reveal deeper neural dynamics, detecting non-linear phenomena overlooked by classical approaches [7, 11, 12]. Such non-linear interactions have been observed in EEG and MEG signals during a range of brain states [13, 14, 15, 16]. This progression toward more advanced computational methods for analyzing brain data highlights the need to better understand the complex nature of brain activity. Advancements in data processing and interpretation uncover new prospects for identifying key neural interactions and hold great promise for developing diagnostic tools for neurological and psychiatric conditions [17, 4].

However, as the complexity of analyzed data properties increases, measuring and identifying statistical relationships between patterns becomes increasingly difficult. This phenomenon is known as the curse of dimensionality. As the dimensionality of observations, understood here as higher order statistical moments across all multiplets of sensors, increases, it becomes exponentially more difficult to find meaningful relationships between variables, making it essential to reduce dimensionality to make statistical models feasible in practice, allowing researchers to focus on the most informative aspects of the data [18, 19]. To address this challenge, a common approach is to reduce the number of variables required to describe the patterns, a process known as dimensionality reduction. This field has seen substantial growth over the past two decades, emerging as a fundamental technique in data analysis. Dimensionality reduction techniques transform datasets with high dimensionality into a lower-dimensional space while retaining the geometry of the data as much as possible, which enables improved classification, visualization, and understanding of the data [20]. These techniques are crucial in high-dimensional settings, as they improve computational efficiency while preserving the essential structure of the data. By focusing on key features, they uncover patterns that are otherwise obscured by noise and high-dimensional variability, facilitating more interpretable models and ultimately leading to more accurate and robust analyses [21, 22, 23, 24].

Recent studies have demonstrated that tensor decomposition methods are particularly effective in handling this multi-dimensionality, yielding robust and interpretable results in neural signal analysis [25]. Tensor factorization techniques enable the capture of patterns across multiple modes - such as time, frequency, and space - offering deeper insights into the underlying dynamics of neural processes [26]. Traditionally, dimensionality reduction has been achieved through techniques like Principal Component Analysis (PCA) and factor analysis, which are designed for matrix (two-dimensional) data. However, to deal with more complex multi-dimensional data, methods such as PARAFAC (Parallel Factor Analysis) generalize factor analysis to decompose tensors into rank-one components, preserving interactions across all modes. Similarly, tensor factorization methods generalize PCA to multi-way data, where the multilinear structure across modes must be preserved. These tensor-based techniques enable more accurate representation and analysis of complex, multi-way data, capturing interactions that would otherwise be lost in traditional two-dimensional methods [22].

The aim of this work is not a dimensionality reduction of raw data themselves but of quantities calculated from them, namely third order statistical moments in the frequency domain. If calculated for all possible index triplets the number of moments is huge even for a single frequency combination. While methods to decompose third order tensors are not new, our key observation is that these methods are mathematically applicable but are physically meaningless for our case as will be explained below. In contrast to those methods our starting point is a physical model, namely that the statistical moments in a relatively large number of sensors were generated from a few sources with a real valued forward mapping and complex valued coupling in source space. We will see that this leads to a different model for the dimensionality reduction, not only having substantially fewer parameters for the same number of components but also allowing those parameters to be used in standard methods for source reconstruction.

## 2 Methods

### 2.1 The Cross-Bispectrum and Its Low Dimensional Model

The cross-bispectral tensor for a set of *N* sensors is defined as:

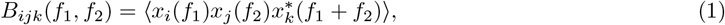

where *x*_*i*_(*f*) is the Fourier transform of the signal at sensor *i*, ⟨·⟩ represents the expectation value, and *f*_1_ and *f*_2_ are the frequencies of interest. This expectation value is estimated by an average over trials or segments.

We assume that the observed signals at the sensors are generated by a small number *M* of latent sources. Let *s*_*m*_(*f*) denote the signal from the *m*-th latent source at frequency *f*, and let *A* ∈ ℝ^*N×M*^ be the mixing matrix that maps the latent sources to the sensors. The observed signal *x*_*i*_(*f*) at sensor *i* can be expressed as a linear combination of the latent source signals:

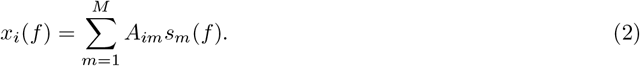

Substituting the mixing model into the definition of the cross-bispectrum, we get:

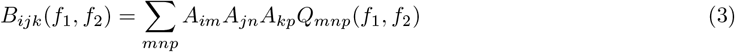

with the definition:

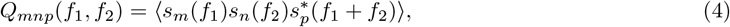

which describes the interactions between the latent sources at the respective frequencies.

The objective of the method is to estimate the mixing matrix *A* and the source cross-bispectrum *Q* by minimizing the difference between the observed bispectrum *B*_*ijk*_ and its modeled form:

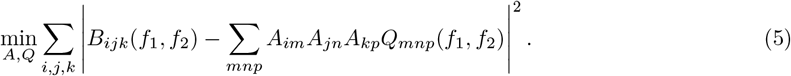

Mathematically, the objective is to solve:

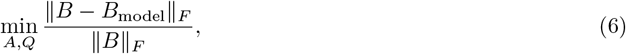

where ∥ · ∥_*F*_ denotes the Frobenius norm, and is used to quantify the relative error between the observed and modeled bispectra.

The optimization problem results in significant dimensionality reduction. For instance, when dealing with *N* = 60 sensors, the original bispectrum tensor *B*_*ijk*_ has *N* ^3^ = 216, 000 complex-valued entries, which is equivalent to 432,000 real-valued observations. However, the number of parameters that need to be estimated in our model is substantially smaller. Assuming *M* = 3 latent sources, the mixing matrix **A** contains 60 × 3 = 180 real-valued entries. The source bispectrum tensor **Q** contains 3^3^ = 27 complex values, or 54 real-valued parameters. Therefore, the total number of parameters to estimate is 180 + 54 = 234. This is a much lower dimensional space compared to the original bispectrum data, offering a significant reduction in complexity while preserving the essential interactions captured by the bispectral analysis.

To fit the model, we use the Levenberg-Marquardt method (LM), which is based on the Jacobian matrix of the forward model. While the method appears to be straightforward, the direct implementation is computationally extremely costly both with regard to computation time and required memory because a Jacobian matrix has size number of observations times number of parameters. On the other hand, if *J* is such a Jacobian matrix and if *x* are all observations stacked to one vector, then LM only requires the knowledge of the quantities *J*^*T*^ *J* and *J*^*x*^, which are typically hundreds of times smaller than *J* itself. We derived rather complicated analytical formulas for *J*^*T*^ *J* and *J*^*T*^ *x*. The concepts of performing the fit are explained in the appendix and the Matlab code for this is provided at [27]. All other code, namely the forward and inverse calculation, visualization, calculation of bispectra, MOCA (see below), and also the example data set can be found in the METH toolbox [28].

### 2.2 Comparison to Other Tensor Decomposition Methods

To contextualize the proposed model, we compare it conceptually to standard tensor decomposition frameworks commonly used in signal processing, namely the Tucker and PARAFAC models. The decomposition method which is closest to our model is the Tucker model. It reads for our case

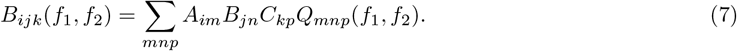

The first difference is that in the Tucker model one has three different ‘mixing matrices’ *A, B* and *C*. The second difference is that for complex valued observations also the mixing matrices *A, B* and *C* can be complex valued. Since the forward mapping is real valued in the standard quasi-static approximation, the ‘mixing matrices’ from the Tucker model cannot be interpreted as topographies of sources in sensor space. In our model, there is only one real valued mixing matrix *A* and only *Q* may be complex valued. A mathematical consequence is that the Tucker model is multi-linear, i.e. assuming e.g. that *B, C* and *Q* are known one can solve for *A* with linear regression. This allows for an iterative search algorithm using only linear regression tools, which is not possible for our model. We emphasize that the Tucker model, while highly flexible, is not well-suited for our application. It is primarily designed for cases where the tensor indices represent different modalities, such as time, frequency, and space. In such cases, assuming identical mixing matrices across all modes would not be appropriate.

A special case of the Tucker model is the PARAFAC model which we mention here for completeness. The PARAFAC (Parallel Factor Analysis) method was introduced independently by Carroll and Chang (1970) [29] and Harshman (1970) [30]. It is a tensor decomposition technique designed to extend matrix factorization methods into higher-order tensors, which allows it to model multi-way data, capturing the latent structures in such data. The main motivation behind PARAFAC is to generalize the notion of decomposition from matrices to tensors, enabling the modeling of multi-dimensional arrays in a way that reflects interactions across all modes. Mathematically, PARAFAC decomposes a tensor into a sum of rank-one tensors. For our third-order tensor, the PARAFAC decomposition is represented as:

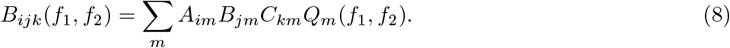

While our expansion can be considered as the Tucker model with additional constraints, this is not the case for the PARAFAC model with the latter assuming for our application that all bispectral couplings between different sources vanish, i.e. *Q*_*mnp*_ vanishes unless *m* = *n* = *p*.

### Minimum Overlap Component Analysis (MOCA)

We recall that our decomposition model is unique up to real valued mixing of the columns of the mixing matrix. To spatially disentangle the source fields contributing to bispectral coupling components, we apply *Minimum Overlap Component Analysis (MOCA)* [31, 32]. MOCA operates on source-space projections obtained by applying a linear inverse operator to sensor-level topographies, and aims to demix overlapping source contributions based on a spatial orthogonality and minimal overlap criterion.

To explain the concept we will assume only two sources. Let *x*_1_, *x*_2_ ∈ ℝ^*M*^ denote two topographies assumed to be mixtures of the topographies of two sources. First, the two topographies are mapped to source space via a linear inverse operator *G*, yielding source estimates

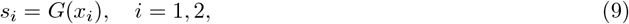

where *s*_*i*_(*m, k*) denotes the value at voxel *m* and direction *k* ∈ {1, 2, 3}. These fields are assumed to be mixtures of the true source fields *q*_*j*_,

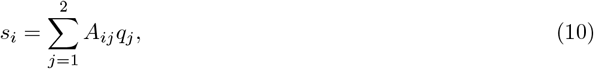

where the unknown fields *q*_1_ and *q*_2_ are assumed to be spatially orthonormal:

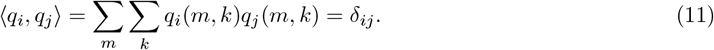

To retrieve these source fields, MOCA first applies a whitening transformation to ensure orthonormality of *s*_1_ and *s*_2_:

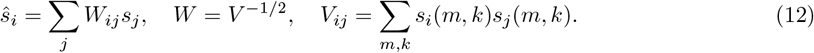

The demixed fields *q*_1_ and *q*_2_ are then obtained by applying a rotation in the whitened 2D subspace:

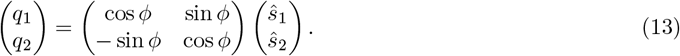

The optimal angle *ϕ* is found by minimizing the spatial overlap between the rotated fields, defined as

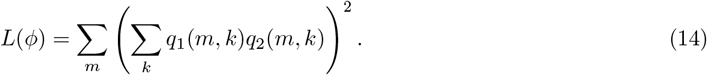

This cost function is minimized analytically via

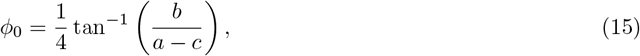

Where

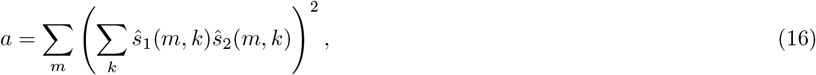

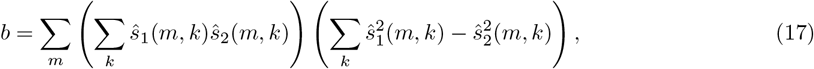

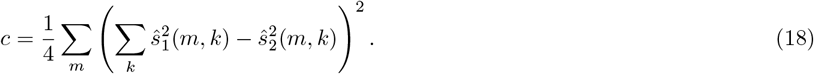

The resulting components *q*_1_ and *q*_2_ represent spatially resolved estimates of the sources contributing to the original bispectral component. MOCA is also generalized to the case of *N >* 2 sources by iteratively applying pairwise rotations that minimize the total overlap cost function [32].

### 2.3 Summary and Methodological Comparison

Figure 1 illustrates the full processing pipeline for our decomposition and analysis framework. The process begins with the **EEG recordings**, which serve as the input signals for analysis. In the first step, we compute the **cross-bispectral tensor**, which is analyzed independently for each frequency pair, focusing on the dominant pair exhibiting maximal bicoherence within the alpha band. This tensor is then subjected to **low-rank decomposition**, which identifies a compact set of latent components that approximate the underlying structure of the data while reducing its dimensionality. These components are subsequently projected into source space via an **inverse mapping**, producing spatial distributions associated with each bispectral component. Since the low-rank decomposition is not unique, MOCA is applied to obtain a unique solution. It does so by assuming that the underlying sources are minimally overlapping in space, resulting in clearly separated source patterns. Finally, the resulting demixed source patterns are subjected to **coupling analysis**, enabling interpretation of cross-frequency interactions in terms of their anatomical origin and functional connectivity. This end-to-end framework is specifically designed to handle nonlinear coupling structures in EEG data and yields interpretable source-level representations of neural interactions.

**Figure 1.**
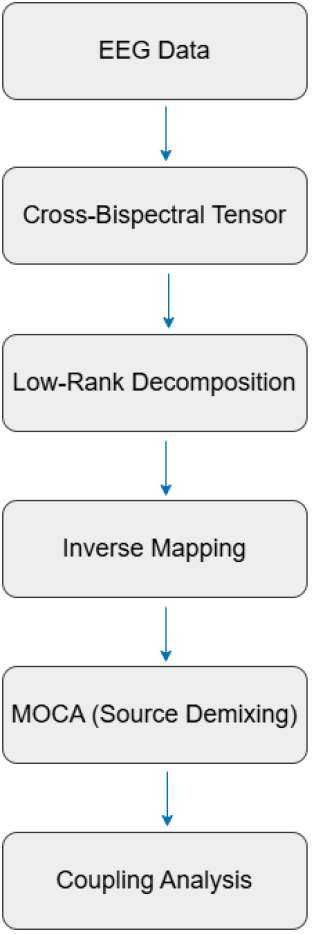
Processing pipeline for the method.

We compare the concept of our method specifically with the Tucker model and, more restrictively, with its special case PARAFAC. All three approaches provide low-rank approximations that preserve the multi-way structure of the bispectral tensor, but they differ in how latent components and interactions are represented. The Tucker model decomposes the tensor into three mode-specific mixing matrices and a core tensor *G*, offering high flexibility but treating the interaction structure in the core as an unconstrained parameter array. In our context, this corresponds to allowing arbitrary couplings between sensor indices without enforcing a generative source model. PARAFAC further constrains the decomposition to a sum of rank-one tensors with shared latent factors across modes; for cross-bispectral data this is equivalent to assuming that *Q*_*mnp*_ vanishes unless *m* = *n* = *p*, i.e. that only self-interactions of sources are present. In contrast, our model uses a single real-valued spatial mixing matrix *A* and a structured source-interaction tensor *Q*_*mnp*_, so that cross-source bispectral interactions are represented explicitly at the level of latent sources. This structure makes the model less flexible than full Tucker but more interpretable and better matched to the generative assumptions underlying cross-bispectral coupling. Because the Tucker model is the most flexible multilinear framework and the closest in form to our decomposition, differing mainly in the structure imposed on the core, it provides the most relevant baseline for comparison.

Table 1 provides a direct comparison between our method and the Tucker model. While Tucker decomposes a tensor into independent mode-specific factors and a core tensor *G*, our method uses a shared spatial mixing matrix *A* and a latent source interaction tensor *Q*, which directly models cross-frequency coupling among latent sources. This makes our approach particularly suitable for capturing non-linear dependencies, while the Tucker model primarily captures linear variance structure. Additionally, the Tucker model allows for distinct rank constraints per mode, whereas our method operates under a single latent dimensionality, yielding a more compact and interpretable representation tailored for interaction modeling in neural signals.

**Table 1:**
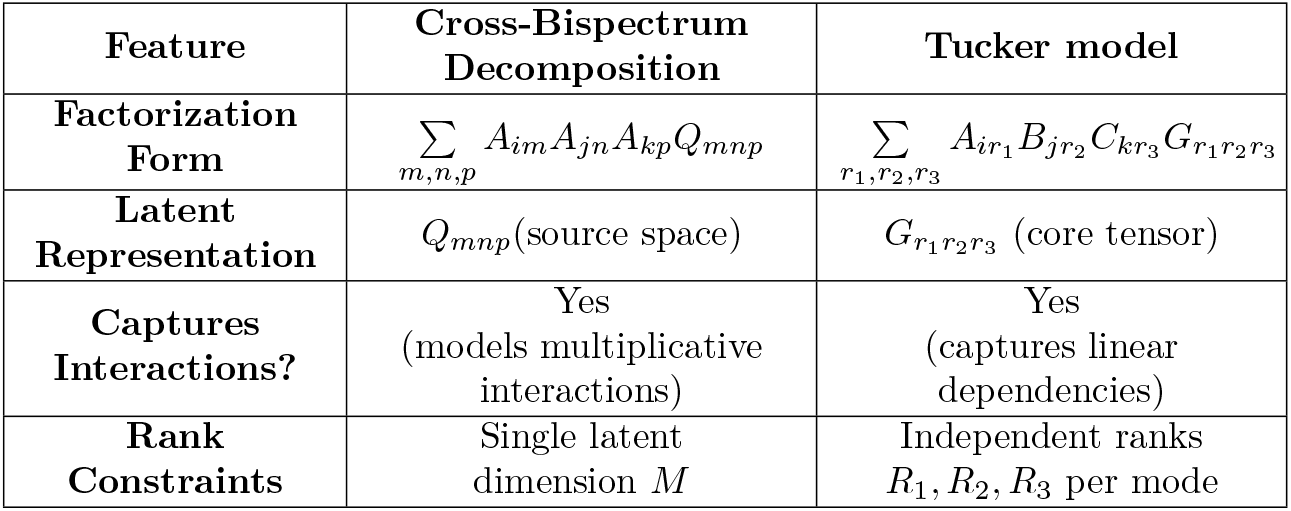
Comparison of the proposed method and the Tucker model.

## 3 Results

### 3.1 Synthesized EEG Data

To assess the performance of the proposed decomposition framework under controlled and interpretable conditions, we developed a simulation pipeline that generates realistic multichannel EEG data using cortical sources. The EEG forward model was computed using the method introduced by Nolte and Dassios [33], which computes the leadfield matrix for a set of dipolar sources distributed throughout a realistic head model. The cortical source space consisted of 5003 candidate dipole locations arranged on a medium-resolution cortical grid embedded in a standard head geometry. Each dipole location was associated with three orthogonal orientation components, allowing arbitrary source orientation. EEG data were generated at 61 simulated scalp electrodes using leadfields provided by the head model, which defines the linear mapping between each source and the scalp electrodes. Simulated source signals were assigned to anatomically plausible cortical locations across both hemispheres, with the number of active sources varied systematically across experiments to evaluate the robustness of the method under different levels of complexity.

To simulate background brain activity, spatially uncorrelated white Gaussian noise was generated across all cortical voxels in source space. For *N* voxels and *T* = 10,000 time points, noise samples were drawn from a standard normal distribution, resulting in a noise matrix of size *T* × *N*. This source-level noise was then forward-projected to the sensor level using the full leadfield matrix, resulting in background activity consistent with volume conduction effects:

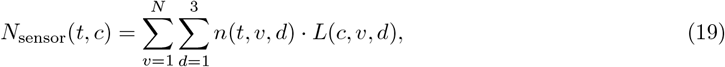

where *n*(*t, v, d*) is the noise at time *t*, voxel *v* in direction *d*, and *L*(*c, v, d*) is the leadfield from voxel *v* to sensor *c* in direction *d*.

The projected sensor-space noise was subsequently band-pass filtered into the alpha (8–12 Hz) and beta (20–30 Hz) bands, and the filtered components were combined with equal weighting to produce background signals with realistic spectral structure. This background activity was then scaled to achieve specific signal-to-noise ratios (SNRs) relative to the signal of interest, enabling systematic evaluation of the method’s robustness to physiologically plausible interference. This procedure ensured physiologically plausible sensor-level noise with spectral content relevant to EEG, while maintaining spatial independence at the source level.

To simulate nonlinear cross-frequency interactions between cortical sources, we constructed coupled signal pairs in which one source was derived through a nonlinear transformation of another, followed by zero-mean normalization and amplitude rescaling. Let *s*_1_(*t*) denote a narrowband oscillatory signal generated by filtering Gaussian white noise within a selected frequency band. Its coupled partner *s*_2_(*t*) was obtained by applying a pointwise quadratic nonlinearity, followed by zero-centering and normalization:

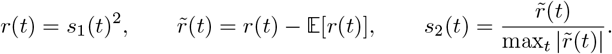

Here, *r*(*t*) is the instantaneous squared amplitude of the original signal *s*_1_(*t*), and 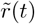 denotes the zero-mean version of *r*(*t*), obtained by subtracting its temporal mean 𝔼 [*r*(*t*)]. The final signal *s*_2_(*t*) is amplitude-normalized to ensure unit dynamic range before being scaled to the desired microvolt level. This transformation creates a nonlinear interaction that introduces energy at the sum frequency *f*_1_ + *f*_2_, which corresponds to 2*f*_1_ in the case of self-squared coupling. The resulting bispectral component reflects second-order phase coupling between *s*_1_ and *s*_2_, reflecting an interaction where energy is redistributed from the base frequency *f*_1_ to its harmonic 2*f*_1_ [7].

Each simulated signal was scaled to an amplitude between 100–300 *µ*V. To ensure that these source-level amplitudes resulted in physiologically realistic EEG signals at the scalp, we evaluated the resulting sensor-level data by computing the maximum and RMS amplitudes across all channels and time points. Across representative simulations, the resulting EEG signals exhibited peak sensor amplitudes in the range of 10– 100 *µ*V, which is consistent with realistic EEG measurements [34, 35]. Higher-amplitude signals modeled dominant sources, while coupled or uncoupled secondary sources were assigned lower amplitudes. Dipole orientations were either fixed or randomly sampled on the unit sphere to account for realistic variability in the direction of current flow. Sensor-level EEG data were generated by forward-projecting the dipolar signals through the leadfield matrix and summing the resulting scalp potentials.

To identify dominant cross-frequency interactions, we first computed the normalized univariate cross-bispectrum across all EEG channels. The bispectral tensor was estimated at each channel using Equation 1. Bicoherence was evaluated over frequency combinations within the alpha band, and the channel with the maximum bicoherence value was selected. The corresponding frequency pair was used to define the target interaction. A full cross-bispectral tensor was then computed across all channels at the selected frequency combination, yielding a complex-valued tensor of size channels × channels × channels. This tensor was subsequently decomposed using the proposed dimensionality reduction method.

This simulation framework provides full control over the spatial, spectral, and coupling properties of the data, enabling rigorous validation of the method under realistic conditions. In the following sections, we report results evaluating the method’s performance across a range of controlled experiments, including variations in source count, dipole orientation, coupling structure, amplitude asymmetry, and noise level. Each scenario was designed to test a specific aspect of the method’s sensitivity, specificity, and robustness.

#### Performance Metrics

To quantitatively evaluate the accuracy of the proposed decomposition framework, we employed the following metrics:

#### Model Error

This metric quantifies the relative reconstruction error between the original cross-bispectral tensor ℬ and its low-rank approximation 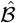. It is defined as:

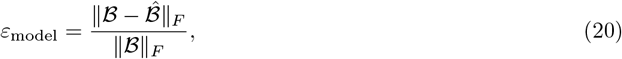

where ∥·∥ _*F*_ denotes the Frobenius norm. Lower values indicate a more accurate representation of the original bispectral structure.

#### Localization Error

Spatial accuracy is assessed by computing the Euclidean distance between the true and estimated source locations for each demixed component:

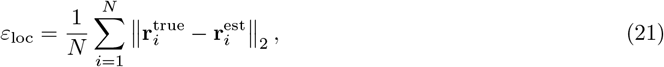

where 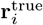 and 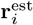 denote the true and estimated peak voxel coordinates, and *N* is the number of sources.

##### 3.1.1 Performance Under Ideal Conditions

To assess the performance of the method under ideal conditions, we simulated two coupled sources in the absence of noise. Source 1 was positioned in the left hemisphere and generated by filtering Gaussian white noise to isolate a 10 Hz signal. Source 2 was placed in the opposite hemisphere and created by applying a pointwise quadratic transformation to Source 1, yielding a signal with dominant energy at 20 Hz. This non-linear transformation established a quadratic phase relationship between the two sources. The transformed signal was centered, normalized to unit range, and scaled to match the amplitude of Source 1 (300 *µ*V). Both sources were projected to the scalp using fixed, pre-defined dipole orientations to ensure spatial reproducibility. No uncoupled sources or background interference were added in this configuration. The results of this simulation are summarized in Fig. 2, which shows the ground-truth dipole configuration (A), a clear bicoherence peak at the expected frequency pair (B), and the true versus demixed source topographies (C).

**Figure 2.**
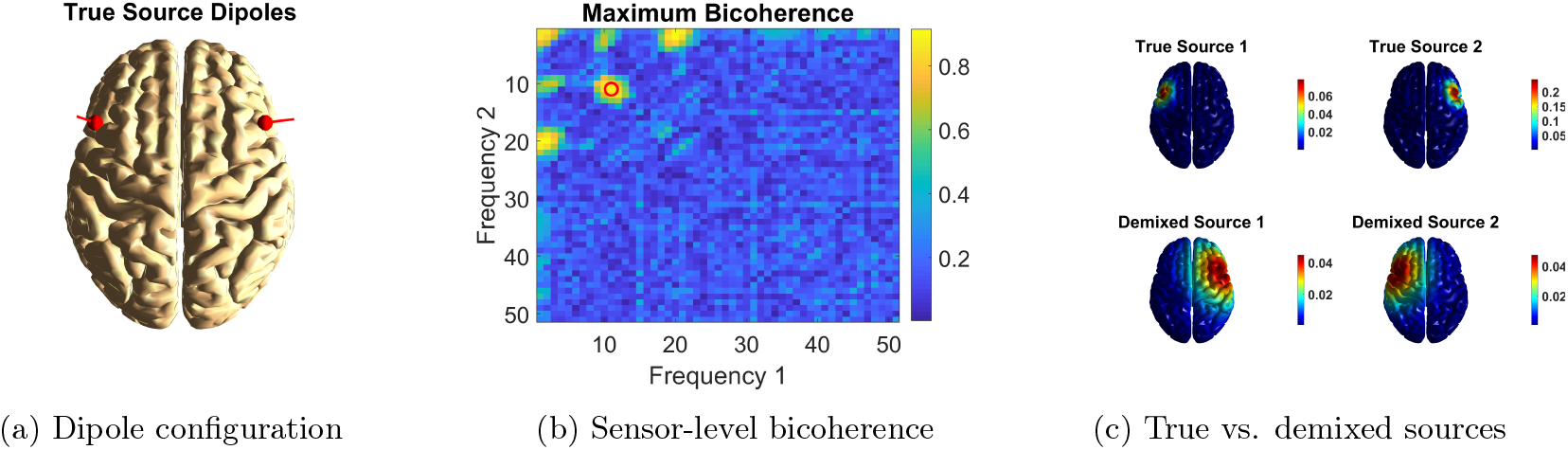
Ideal simulation results. (A) True dipole locations and orientations of the two coupled sources. (B) Bicoherence map at the dominant sensor, showing a peak at bin (11, 11), corresponding to 10–10 Hz interaction. (C) True and recovered source maps from MOCA decomposition, confirming accurate localization. **Note:** Frequency bin *f* corresponds to (*f* − 1) Hz. Here, *f*_1_ = *f*_2_ = 10 Hz and *f*_3_ = *f*_1_ + *f*_2_ = 20 Hz.

##### 3.1.2 Robustness to Signal-to-Noise Ratio

To assess the robustness of the proposed model under varying noise conditions, we conducted a controlled simulation study involving two interacting cortical sources. The source dynamics were designed to generate a cross-frequency coupling signature and both signals were amplitude-normalized and scaled to 300 *µ*V, ensuring comparability across conditions. Dipole orientations were fixed and orthogonal to prevent degeneracy in the forward model and facilitate interpretable topographies. This setup provided a consistent ground truth for evaluating performance and algorithmic stability. Noise was scaled per trial to achieve a defined signal-to-noise ratio (SNR) using the power ratio of the signal to the background activity. The SNR in decibels was computed using the standard logarithmic formula:

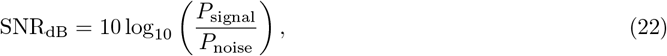

SNR was systematically varied from − 15 dB to +15 dB in 1 dB increments. For each SNR level, ten independent trials were conducted with newly sampled randomized noise. Two performance metrics were computed per trial: (1) the model reconstruction error, and (2) the source localization error. Figure 3 summarizes the results. In the top panel, the model error remains relatively high between SNRs of –15 dB to –10 dB and then decreases smoothly as SNR increases. Within this regime, the cross-frequency coupling signal is heavily masked by background activity, and the model struggles to fully reconstruct the bispectral structure. Beginning around –7 dB, the error decreases steadily and continues to drop across the entire SNR range, reaching values below 0.1 by +15 dB. This marks the transition point at which the bispectral structure becomes reliably detectable. At higher SNRs, variability becomes minimal and the error settles into a smooth decline, reflecting reliable reconstruction of the bispectral structure. The error bars in this range further reinforce stability: while variability is substantial at very low SNR (*<* − 10 dB), it contracts quickly as SNR increases and becomes minimal from roughly –5 dB onward, suggesting that once the coupling signal exceeds the noise floor, the model’s performance becomes robust across runs. The bottom panel illustrates localization error, which declines more steeply and with less fluctuation. Localization error begins around 1–2 mm at the lowest SNRs and decreases rapidly between –7 dB and –3 dB, reaching 0 mm with no variance for SNRs ≥ 0 dB.

**Figure 3.**
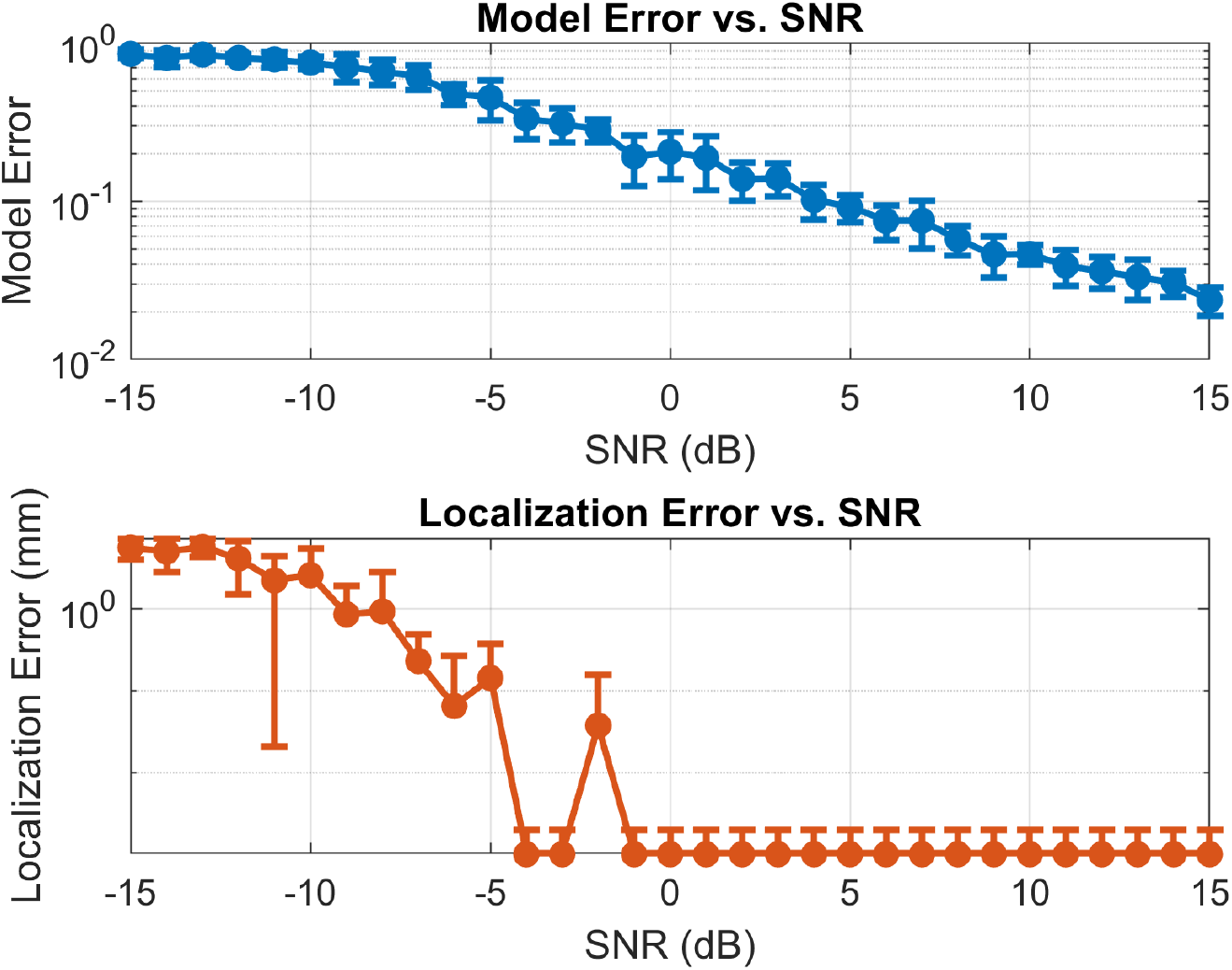
Robustness of the method across signal-to-noise ratios (SNR). Top: model reconstruction error; bottom: source localization error after MOCA demixing. Each data point represents the mean across 10 simulation trials; error bars indicate standard deviation.

These results demonstrate the method’s sensitivity to low SNR, as expected given the higher-order nature of bispectral coupling. Importantly, they also establish its robustness and reliability in moderate-to-high SNR regimes, which are representative of typical empirical EEG recordings [36, 37, 38]. The observed transition region around 0 dB delineates the practical lower bound for successful source-level bispectral decomposition and highlights the effectiveness of the proposed framework in realistic conditions.

##### 3.1.3 Influence of Coupling and Dipole Orientation

We systematically evaluated the influence of cross-frequency coupling and dipole orientation on source separation performance under background noise. Each condition involved two sources located in distinct cortical regions, with either parallel or orthogonal dipole orientations (see Fig. 4). In the uncoupled configurations, sources oscillated independently at 10 and 12 Hz, while in the coupled conditions they were quadratically coupled at 10 Hz. To isolate the effects of geometric and coupling structure, the SNR was held constant at 0 dB, a level previously identified as sufficient for reliable reconstruction and accurate source localization. Across all cases, perfect localization accuracy (0.00 mm error) was achieved, but the model (bispectral) error varied depending on the presence of coupling and the dipole geometry. Specifically, the highest model error occurred in the uncoupled conditions consistent with the lack of cross-frequency interaction between sources. A summary of bispectral decomposition error across all tested conditions is provided in Table 2.

**Table 2:** Bispectral decomposition error (bsfit) across conditions.

| Condition | Dipole Orientation | Model Error (bsfit) |
| --- | --- | --- |
| Uncoupled | Parallel | 0.562 |
| Coupled | Parallel | 0.284 |
| Uncoupled | Orthogonal | 0.672 |
| Coupled | Orthogonal | 0.208 |

**Figure 4.**
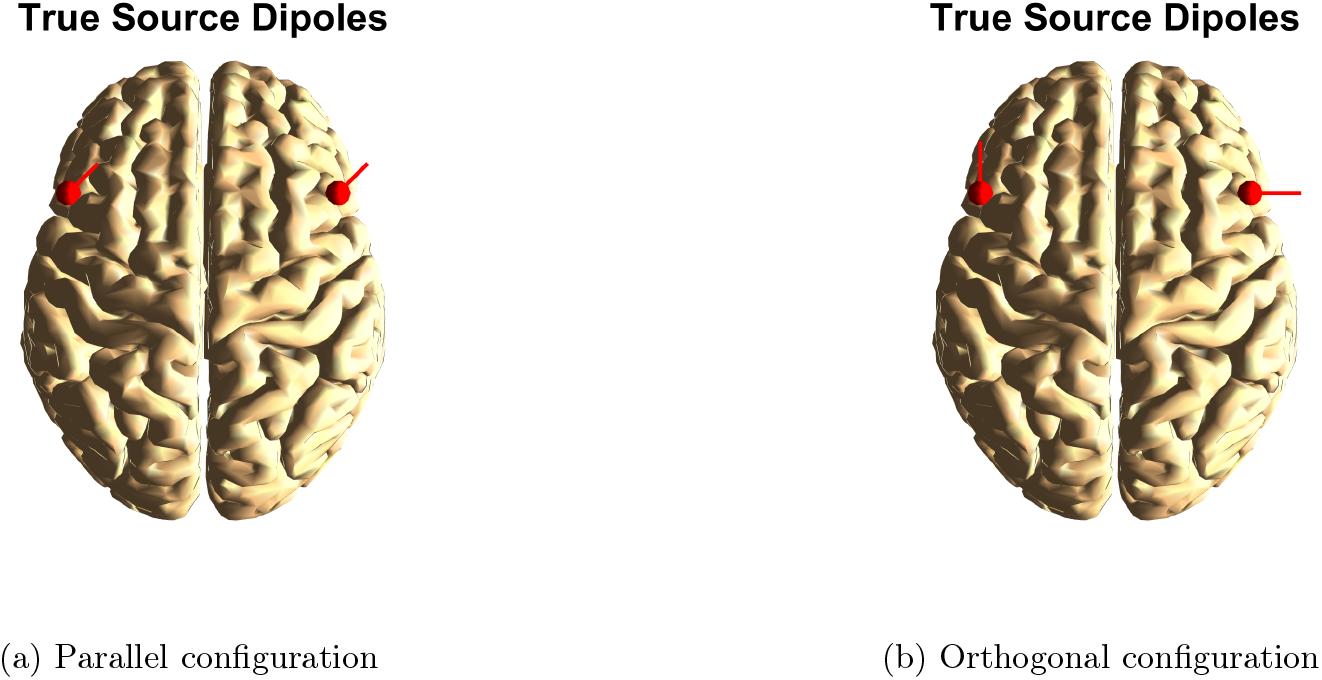
Dipole orientations used in the simulations.

To further examine the influence of dipole orientation variability on the model’s performance, we conducted 100 simulations with randomized dipole directions under the coupled condition. In each run, two dipoles were placed at fixed cortical locations and injected with quadratically coupled signals at 11 Hz. While dipole positions and signal amplitudes remained constant, the orientation vectors were sampled randomly from the unit sphere to mimic realistic variability in source geometry. This design isolates the effect of directionality from spatial location or coupling strength, allowing us to assess how misalignment of dipole vectors influences model estimation. All simulations preserved perfect localization accuracy via MOCA (0.00 mm), confirming spatial robustness. However, the model error exhibited moderate variability, with most values falling between approximately 0.15 and 0.30, alongside a few higher-error outliers reaching up to 0.82, as shown in Fig. 5. The histogram illustrates the distribution of bispectral model errors obtained across 100 independent simulations, each with randomly oriented dipole directions, thereby quantifying the variability introduced by source geometry alone. The distribution is strongly concentrated in the lower error range, indicating that the majority of random orientations yield similarly accurate fits. These results highlight that while coupling is consistently detected, the quality of the fit remains influenced by source orientation, with certain geometric configurations producing noticeably higher errors. This dependence likely reflects how orientation affects source mixing in sensor space, shaping how distinct or overlapping the bispectral features appear across channels. Thus, evaluating decomposition performance across a distribution of geometries provides critical insight into the model’s generalizability under realistic conditions.

**Figure 5.**
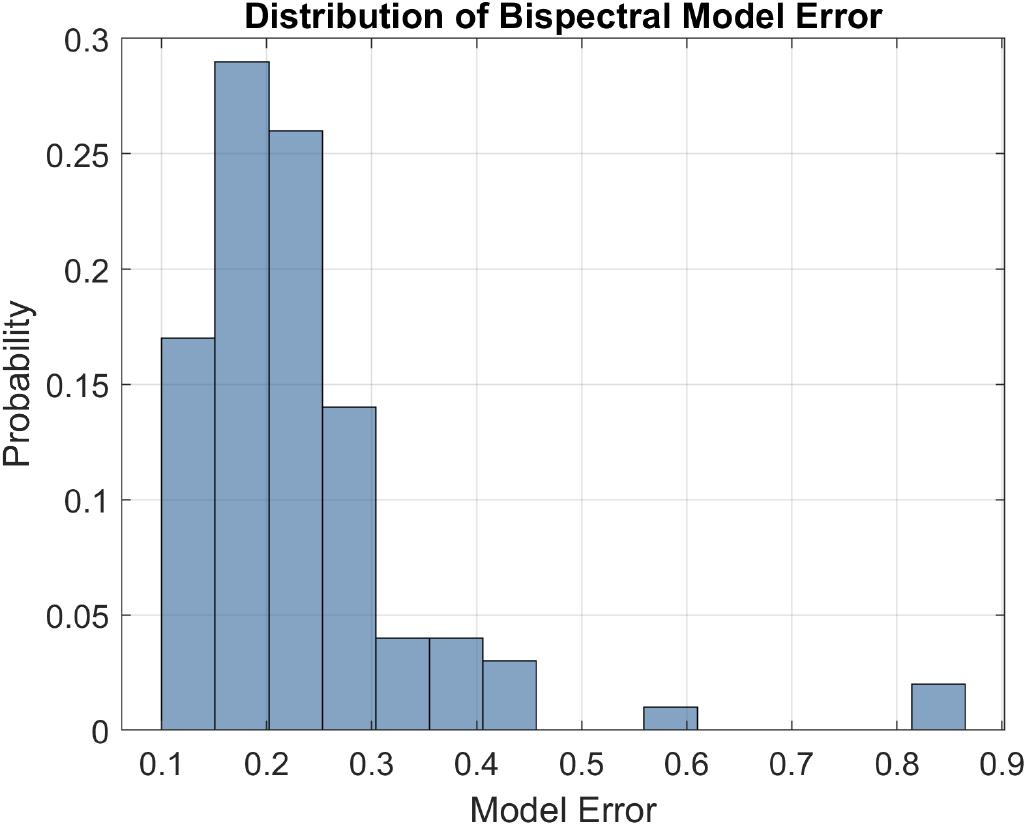
Histogram of bispectral model errors across 100 simulations with randomized dipole orientations (coupled condition).

##### 3.1.4 Performance with Increased Complexity

We next tested the method in a more complex and challenging setting involving four sources. All sources were placed in distinct locations across both hemispheres. The corresponding dipole visualization (Fig. 6) illustrates the spatial layout and dipole orientation of all four sources, providing a clear reference for the anatomical configuration used in the simulation. We simulated six scenarios with varying configurations of coupling and amplitude imbalance with SNR fixed at 0 dB.

**Figure 6.**
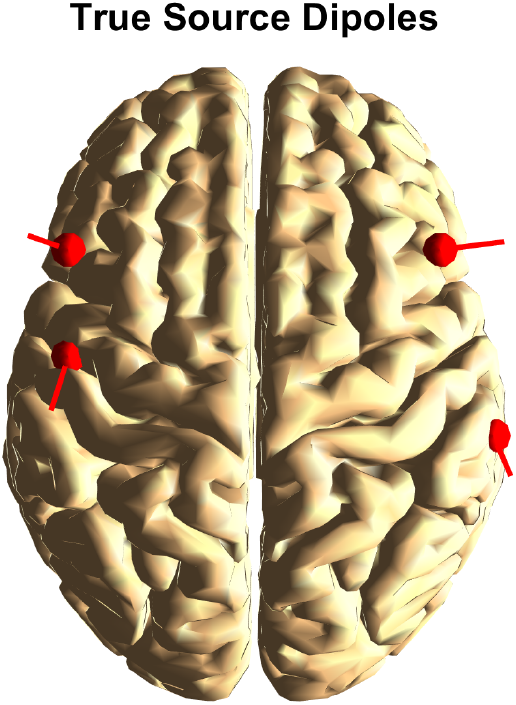
Visualization of true source dipoles. Red arrows indicate the orientation and location of the four simulated dipoles on the cortical surface.

The results presented in Table 3 reveal that the performance of the model is governed by several interacting factors: the presence of true coupling, the relative amplitudes of coupled versus uncoupled sources, and the overall consistency of the signal. A consistent pattern emerges in conditions where the coupled sources dominate in amplitude, either because all sources are coupled (Case 6) or because the uncoupled sources are comparatively weak (Cases 2 and 4). In these settings, the model achieves lower decomposition error, suggesting that when the dominant contributors to the sensor-level signal reflect coherent cross-frequency interactions, the low-rank bispectral approximation provides an accurate and compact representation. In contrast, when uncoupled sources are energetically comparable to the coupled ones (as in Cases 1 and 3), the model exhibits moderately higher errors compared to the amplitude-imbalanced cases. In these cases, the uncoupled components contribute non-interacting spectral energy that is not well-explained by a low-rank factorization of the cross-bispectrum, effectively acting as structured noise relative to the modeled interaction. The highest error is observed in the null condition (Case 5), where all sources are uncoupled despite having strong amplitudes. Here, the model fails not because of algorithmic instability, but because there is no coherent bispectral structure to be captured. This serves as an important negative control, confirming that the decomposition does not artificially impose coupling structure when it is absent. Importantly, the performance does not appear to be tied to the specific frequency of interaction (e.g., 10 Hz vs. 12 Hz). Rather, it is the consistency and prominence of the bispectral interaction relative to other signal components that governs model accuracy.

**Table 3:** Summary of all simulation cases with coupling configurations, source amplitudes, and bispectral decomposition error.

| Case | Coupling Configuration | Amplitudes ($\mu\text{V}$) | Model Error (bsfit) |
| --- | --- | --- | --- |
| 1 | S1–S2 coupled at 10 Hz; S3=12 Hz and S4=9 Hz uncoupled | All sources: 300 | 0.357 |
| 2 | S1–S2 coupled at 10 Hz; S3=12 Hz and S4=9 Hz uncoupled | Coupled: 300, Uncoupled: 100 | 0.285 |
| 3 | S3–S4 coupled at 12 Hz; S1=10 Hz and S2=11 Hz uncoupled | All sources: 300 | 0.304 |
| 4 | S3–S4 coupled at 12 Hz; S1=10 Hz and S2=11 Hz uncoupled | Coupled: 300, Uncoupled: 100 | 0.208 |
| 5 | All sources uncoupled (S1=10 Hz, S2=11 Hz, S3=12 Hz, S4=12 Hz) | All sources: 300 | 0.894 |
| 6 | All sources coupled: S1=10 Hz, S2=10 Hz, S3=12 Hz, S4=12 Hz | S1/S2: 100, S3/S4: 300 | 0.125 |

Figure 7 shows representative spectral and spatial outcomes from two types of coupling scenarios. The left column corresponds to conditions where 10 Hz coupling is dominant (as in Cases 1 and 2), while the right column illustrates scenarios with dominant 12 Hz coupling (as in Cases 3 and 4). In both examples, the bicoherence matrices clearly highlight the expected frequency pair, indicating that the model reliably detects quadratic phase interactions despite the presence of additional uncoupled sources. The lower panels compare the true and demixed source distributions, demonstrating accurate spatial localization of the coupled sources. Importantly, this accuracy is maintained even in the presence of amplitude imbalance, where the coupled sources may be either stronger or weaker relative to uncoupled components. Overall, these findings are particularly significant given the presence of multiple active sources, mixed coupling states, and amplitude asymmetries. The error metric, computed as the normalized difference between the original bispectrum and the reconstructed low-rank model, directly reflects the fidelity of the factorization. A low value indicates that the bispectral structure captured by the model accounts for the dominant interaction features present in the data, with minimal residual variance. In this context, the low-rank constraint serves not only to reduce dimensionality but to impose interpretability and robustness in the presence of noise and irrelevant spectral features.

**Figure 7.**
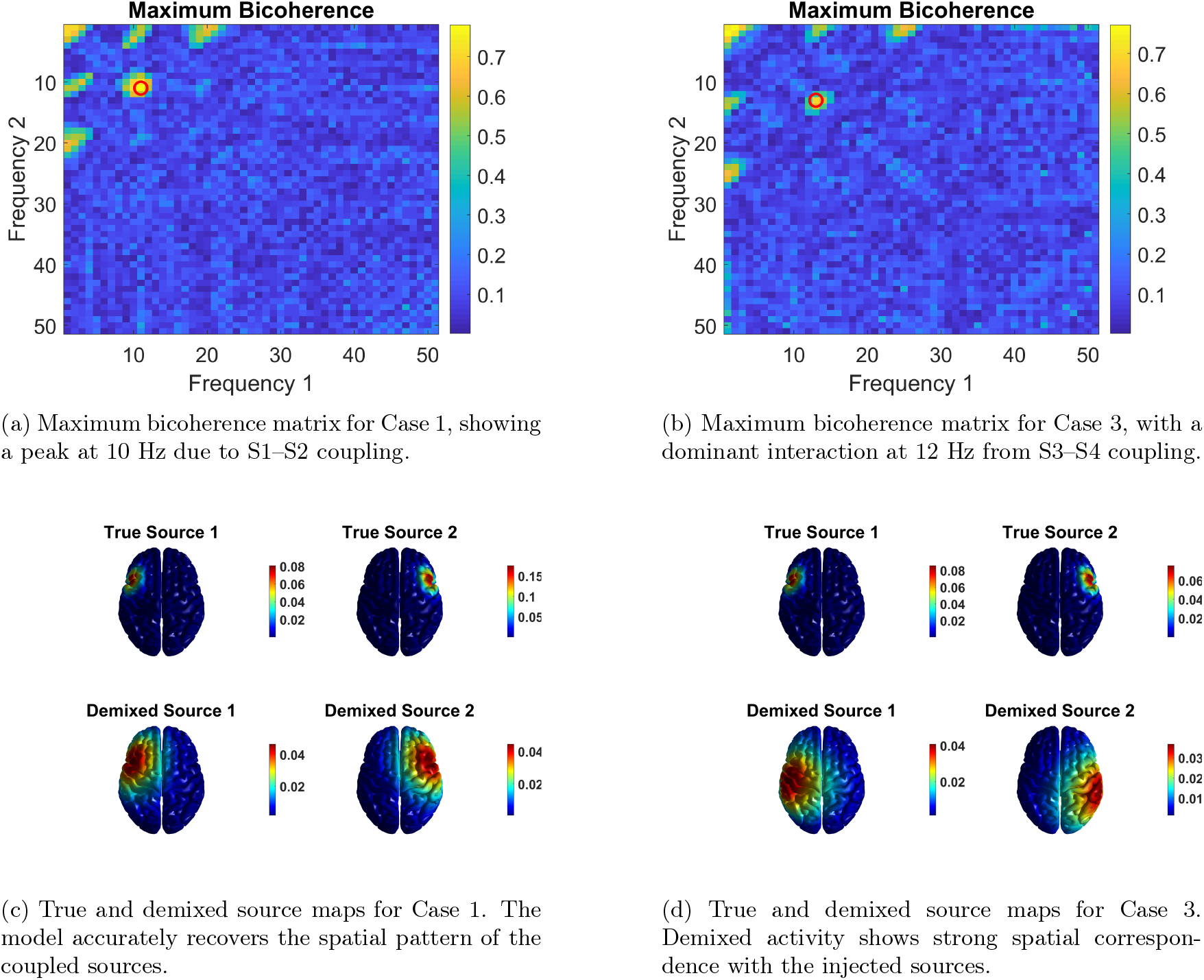
Spectral and spatial decomposition results for representative cases. The left involves 10 Hz coupling between S1 and S2 with additional uncoupled sources, while the right features coupling at 12 Hz between S3 and S4. Top row: sensor-level bicoherence maps with red circles marking the frequency pair of maximum coupling. Bottom row: comparison between true cortical sources and demixed components after bispectral decomposition. **Note:** Frequency bin *f* corresponds to (*f* − 1) Hz.

### 3.2 Application to Real EEG Data

To examine the performance of our approach on empirical data, we applied it to resting-state EEG recordings from 24 neurologically healthy individuals. EEG was acquired using a 64-channel system with Ag/AgCl electrodes positioned according to the international 10–20 layout, with a sampling rate of 1 kHz. Participants rested in an eyes-closed condition for 5 to 10 minutes while continuous EEG was recorded. Preprocessing included artifact correction using independent component analysis (ICA), followed by band-pass filtering between 0.1 and 70 Hz, downsampling to 256 Hz, and re-referencing to the common average. For analysis, the data were segmented into non-overlapping 2-second epochs (512 samples each). The dataset was provided by the Department of Psychiatry at the University Medical Center Hamburg-Eppendorf. Further details regarding the recording protocol and preprocessing can be found in [39, 40]. This dataset allowed us to examine cross-frequency interactions in the absence of external stimulation, offering an optimal testbed to investigate cross-frequency interactions in an undisturbed resting state.

Bicoherence was computed across all sensors and alpha-band frequency combinations (8–12 Hz). As shown in Fig. 8A, the resulting channel-wise coupling matrices exhibited spatially structured variation, with highest values over posterior electrodes. This analysis revealed a clear peak at (13, 13), corresponding to a coupling between 13 Hz and 13 Hz (i.e., *f*_1_ = *f*_2_ = 13 Hz), yielding a summed interaction at *f*_3_ = *f*_1_ + *f*_2_ = 26 Hz (Fig. 8B). Based on this peak, we constructed the cross-bispectral tensor at the corresponding frequency pair and applied a rank-2 decomposition to extract the dominant coupling structure. To localize the spatial origin of the coupled components, the resulting spatial modes were projected to source space using an eLORETA-based inverse operator. The resulting mixed sources (Fig. 9A) exhibited bilateral posterior patterns with substantial spatial overlap. We then applied MOCA to unmix these overlapping components, yielding two spatially distinct sources in left and right parietal cortex (Fig. 9B). This result demonstrates the method’s capacity to extract interpretable and anatomically plausible generators of bispectral coupling from EEG recordings.

**Figure 8.**
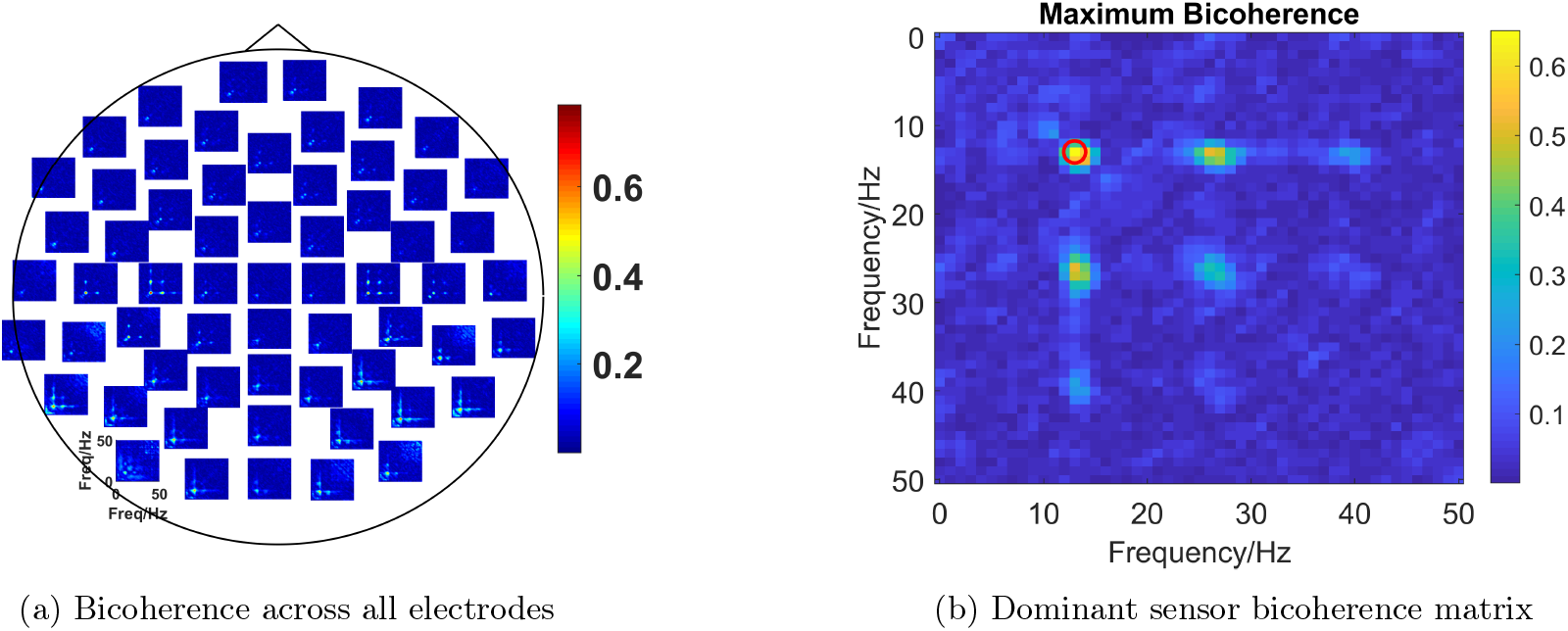
Sensor-level analysis for one subject. (A) Bicoherence across all channels; each square represents the *f*_1_ × *f*_2_ coupling matrix for a single EEG sensor. (B) Bicoherence matrix at the dominant channel (index 15), showing a peak at (13,13), corresponding to a 13–13 Hz interaction.

**Figure 9.**
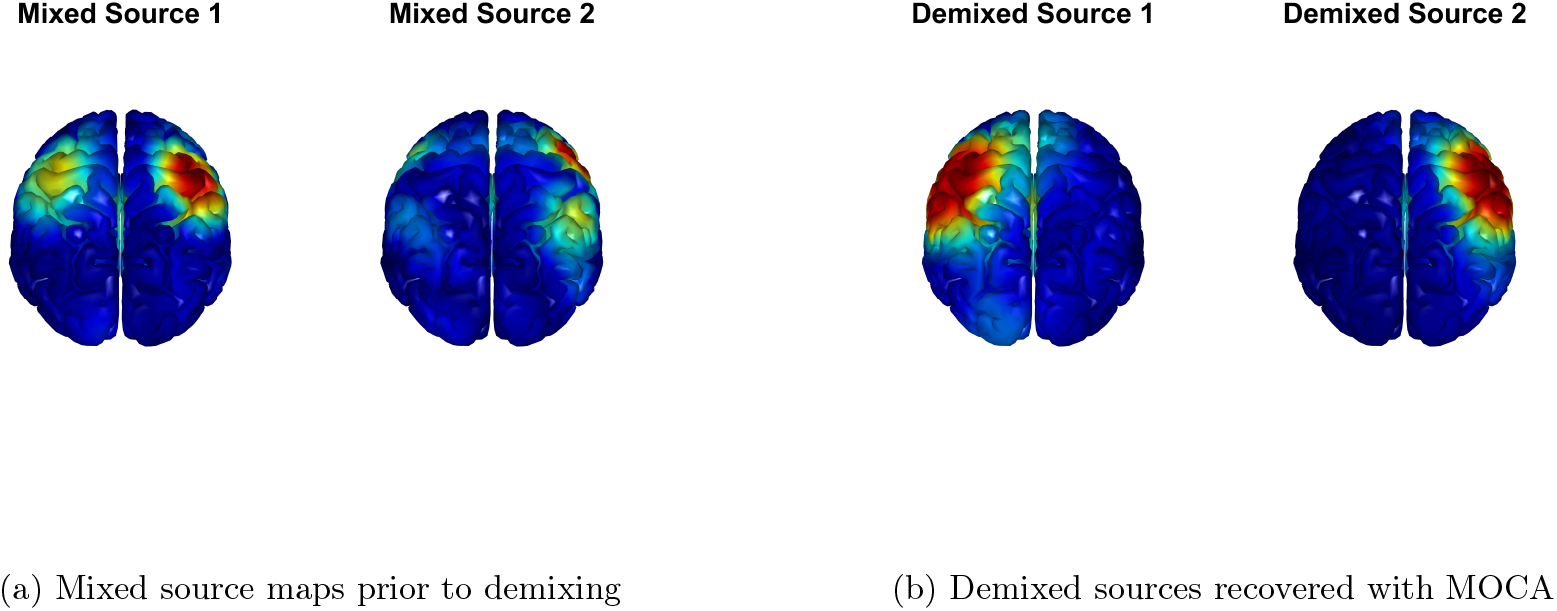
Source-level reconstruction from bispectral components. (A) Mixed spatial components projected via eLORETA show bilateral activation. (B) After demixing with MOCA, two distinct sources emerge in left and right parietal cortex.

## 4 Discussion and Conclusion

We introduced a low-rank decomposition framework tailored to EEG data, enabling the extraction of spatially and spectrally resolved source-level representations of nonlinear neural interactions. The method directly factorizes the observed third-order bispectral tensor into a structured generative model consisting of a spatial mixing matrix and a source-level interaction tensor. This contrasts with standard tensor decomposition methods, such as the Tucker model and its special case PARAFAC, which either assume multilinear interactions or prioritize variance preservation without modeling coupling structure explicitly. A key advantage of the proposed model lies in its substantial reduction in parameter complexity compared to unconstrained bispectral representations. A full cross-bispectral tensor scales cubically with the number of channels, requiring *N* ^3^ complex parameters per frequency pair, where *N* is the number of sensors. In contrast, our model expresses the bispectrum as a structured product of a mixing matrix and a low-dimensional source interaction tensor, reducing the parameter space to *NM* + 2*M* ^3^ real valued parameters where *M* ≪ *N* is the number of latent sources. This not only improves computational efficiency but also enhances model interpretability. By enforcing this low-rank structure, the method effectively suppresses spurious high-dimensional variability and focuses on the dominant nonlinear interactions, which is particularly beneficial in noisy or high-dimensional EEG data. The reduced parameter space further enables the application of nonlinear optimization methods without overfitting, even in the presence of limited or noisy data.

Systematic simulations demonstrated that the method robustly identifies quadratically coupled sources under a range of realistic conditions, including variable signal-to-noise ratios (SNR), amplitude imbalances, dipole orientation variability, and the presence of multiple interacting and non-interacting sources. Importantly, we observed that the model error varied in a principled manner across conditions: it remained low when true cross-frequency coupling was present and dominant, and increased appropriately in the absence of coupling or when uncoupled sources dominated the signal. This behavior indicates that the method does not trivially impose structure but rather reflects the underlying interaction landscape of the data. Localization error remained near zero across nearly all simulation settings, including those with randomized dipole orientations, confirming that the spatial mapping of demixed components is highly robust. This dissociation between model reconstruction error and localization accuracy suggests that while the quality of the low-rank approximation may be sensitive to signal geometry, the spatial extraction of dominant coupled sources remains stable, a favorable property for applications requiring anatomical interpretability.

In comparison to classical tensor models, our method imposes a more constrained but interpretable model structure, where a single latent dimension governs source interactions and spatial mixing. While the Tucker model affords flexibility through independent mode-wise projections, it does not preserve the interaction-specific structure of the cross-bispectrum and lacks a natural mapping to source space. In contrast, our approach yields a decomposition that is both spectrally and anatomically interpretable, facilitating analysis of source coupling.

The model fit is not unique. Similar to a PCA analysis of a covariance matrix, the topographies of the first *M* components are only up to a mixing of these components. The result is then the subspace spanned by these components and not the topographies of the individual sources. We here chose MOCA to demix these components, but many other subspace methods exist which could also be applied to localize the sources from the subspace [41, 42, 43, 44, 45].

Application to real EEG recordings demonstrated that the method captures physiologically plausible patterns of nonlinear synchronization, consistent with prior observations of alpha-band coupling during rest. The recovered source components exhibited distinct spatial distributions after applying MOCA, which resolves the non-uniqueness of the decomposition by assuming that true sources overlap minimally in space. This assumption yields unique and anatomically interpretable source patterns, and the residual error remained low, supporting the adequacy of the low-rank bispectral model in empirical data.

Overall, the proposed method provides a principled approach to extracting interpretable cross-frequency interactions from EEG data. By combining low-rank tensor modeling, nonlinear optimization, and spatial demixing, it addresses critical limitations of existing techniques and offers a tractable means of studying higher-order coupling phenomena in large-scale electrophysiological datasets. Future work may extend the framework to other forms of coupling, incorporate temporal dynamics, or adapt it for group-level analysis across participants.

## Acknowledgments

This work was supported by the European Research Council, project cICMs, ERC-2022-AdG-101097402 (awarded to A.K.E.). Views and opinions expressed in this paper are those of the authors only and do not necessarily reflect those of the European Union or the European Research Council. Neither the European Union nor the granting authority can be held responsible for them. This research was also partially funded by the German Research Foundation (DFG, TRR169/B1/B4).

## Appendix

In this appendix we sketch technical details how to perform the model fit. We first recall the model for an empirical complex valued cross spectral tensor *B*_*ijk*_ at a triple of sensors with indices *i, j, k* calculated at some frequencies:

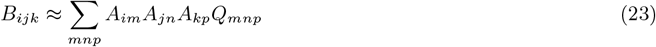

*A* is a real valued matrix and *Q* a complex valued tensor. The indices *m, n, p* run from 1 to *M* for *M* sources which is assumed to be much lower than the number of sensors *N*.

We apply the Levenberg-Marquardt (LM) algorithm to minimize the cost function

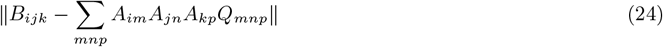

where ∥ · ∥ denotes Frobenius norm, i.e. the absolute values of all elements are squared and then summed.

The LM algorithm is an iteration which first of all requires an initial guess. Such an iteration is not necessary if the model is exactly true and we will use this to construct the initial guess. We first concatenate two out of the three indices of *B* into a single index, say *r*, running from 1 to *N* ^2^ combining the indices *i* and *j*. Then, assuming that the model is correct, (23) has the form

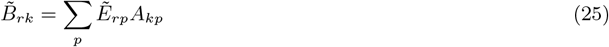

Where 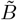 is the reshaped version of the tensor *B*, and 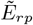 are the reshaped elements of the matrix constructed from

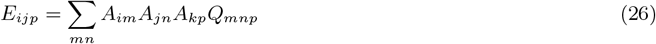

In matrix form this reads

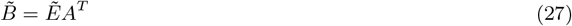

Separating real and imaginary parts, denoted as 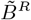 and 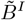, and analogously for 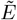, we get

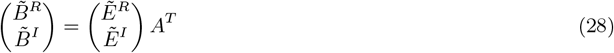

Since *A*^*T*^ is of size *M* × *N* and *M < N* the matrix on the left side of (28) has (at most) rank *M*. We recall that this is the case if the model is exactly true. For empirical data, we make a low rank approximation, and we approximate *A* by the first *M* singular vectors of the matrix

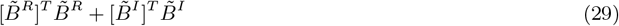

Note that *A* is only defined up to a transformation within source space using any transformation matrix *T* of size *M* × *M*, because it can be absorbed completely by redefining *Q*. It is therefore sufficient to find the space spanned by the topographies of the columns of *A*.

If *A* is given, we are left with a quadratic minimization problem to find *Q*. That can be solved in closed form in a single step. However, since we are dealing with third order tensors, the Matlab code is extremely tedious, and we refer the reader to this code available at [27] for the details. We illustrated the procedure for concatenating the first two indices of *B*_*ijk*_ but it applies equally for any combination of two indices. We test all three possibilities and take that *A* with lowest fit error.

With this initial value for *A* and *Q* we apply the LM algorithm. We here only recall the basic principle. Say, we have some empirical values, say a vector *y*, and a model *f* (*x*) with model parameters *x*, and we want to minimize the cost function *L*(*x*) = ∥*y* − *f* (*x*)∥. At the *k*th step we have an estimated solution 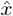, which we want to improve. Then we first linearize *f* around 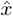 writing 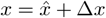 and

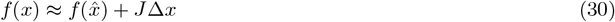

where *J* is the Jacobian matrix, i.e.

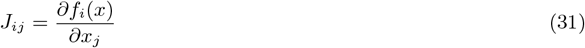

calculated at 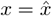. The approximate cost function

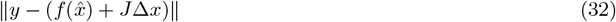

can be minimized in one step with the solution

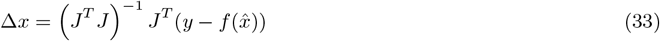

To minimize the full cost function, a regularization is introduced, and a trial estimate is calculated as

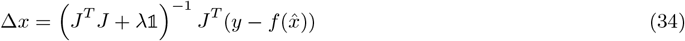

where 𝟙 is the *P* × *P* identity matrix for *P* parameters, and *λ* is a regularization parameter with an essentially arbitrary initial value. If the trial estimate 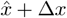 lowers the cost function, this trial estimator is taken as the new estimator and *λ* is decreased by some factor. If the cost function increases, the estimated solution is not changed and *λ* is increased.

Since the model is a nonlinear but simple function of the parameters, the calculation of *J* is straight forward, although tedious. A crucial observation is that we never need *J* itself, but only the matrix *J*^*T*^ *J* and the vector 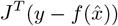, both of which are tiny compared to *J* itself. However, the coding of that is even more tedious than the coding for *J*, and we (again) refer the reader to [27] for our implementation. [28]

For 61 channels and for order *M* = 3 sources the algorithm needs about 300 ms on an ordinary laptop to find the minimum, for *M* = 5 it is around 1 s, and for *M* = 10 around 10 seconds. With the initial estimate for *A* as constructed above the fit error is already remarkably close to the final error. E.g. for *M* = 3 and *f* = 10 Hz (which is coupled to 30 Hz) the final relative fit error is 0.37 whereas the fit error for the initial estimate is 0.40. We get similar results across all frequencies and model orders. In contrast, starting with a random initial guess for *A* the corresponding relative fit error is around 0.9999 with final results usually identical, although with a random initial guess the algorithm sometimes gets trapped in a local minimum.

## Notes

### Competing Interest Statement

The authors have declared no competing interest.

## References

[1] Fumikazu Miwakeichi, Eduardo Martínez-Montes, Pedro A. Valdes-Sosa, Nobuaki Nishiyama, Hiroaki Mizuhara, and Yoko Yamaguchi. Decomposing EEG data into space–time–frequency components using parallel factor analysis. NeuroImage, 22(3):1035–1045, 2004. doi: 10.1016/j.neuroimage.2004.03.039.

[2] Erol Başar. The theory of the whole-brain-work. International Journal of Psychophysiology, 60(2):133–138, 2006. doi: 10.1016/j.ijpsycho.2005.12.007.

[3] Michel Le Van Quyen and Anatol Bragin. Analysis of dynamic brain oscillations: Methodological advances. Trends in Neurosciences, 30(7):365–373, 2007. doi: 10.1016/j.tins.2007.05.006.

[4] Guido Nolte and Klaus-Robert Müller. Localizing and estimating causal relations of interacting brain rhythms. Frontiers in Human Neuroscience, 4:209, 2010. doi: 10.3389/fnhum.2010.00209.

[5] Viktor K. Jirsa and Viktor Müller. Cross-frequency coupling in real and virtual brain networks. Frontiers in Computational Neuroscience, 7:78, 2013. doi: 10.3389/fncom.2013.00078.

[6] Tamanna T. K. Munia and Selin Aviyente. Multivariate analysis of bivariate phase-amplitude coupling in EEG data using tensor robust pca. IEEE Transactions on Neural Systems and Rehabilitation Engineering, 29: 1268–1279, 2021. doi: 10.1109/TNSRE.2021.3092890.

[7] Forooz Shahbazi Avarvand, Sarah Bartz, Christina Andreou, Wojciech Samek, Gregor Leicht, Christoph Mulert, Andreas K. Engel, and Guido Nolte. Localizing bicoherence from EEG and MEG. NeuroImage, 174: 352–363, 2018. doi: 10.1016/j.neuroimage.2018.01.044.

[8] Elizabeth Heinrichs-Graham and Tony W. Wilson. Presence of strong harmonics during visual entrainment: A magnetoencephalography study. Biological Psychology, 91(1):59–64, 2012. doi: 10.1016/j.biopsycho.2012.04.008.

[9] David Sherman, Ning Zhang, Shikha Garg, Nitish V. Thakor, Marek A. Mirski, Mirinda Anderson White, and Melvin J. Hinich. Detection of nonlinear interactions of EEG alpha waves in the brain by a new coherence measure and its application to epilepsy and anti-epileptic drug therapy. International Journal of Neural Systems, 21(2):115–126, 2011. doi: 10.1142/S0129065711002754.

[10] Felix Darvas, Jeffrey G. Ojemann, and Lars B. Sorensen. Bi-phase locking—a tool for probing non-linear interaction in the human brain. NeuroImage, 46(1):123–132, 2009. doi: 10.1016/j.neuroimage.2009.01.034.

[11] Federico Chella, Vittorio Pizzella, Filippo Zappasodi, Guido Nolte, and Laura Marzetti. Bispectral pairwise interacting source analysis for identifying systems of cross-frequency interacting brain sources. Physical Review E, 93(5):052420, 2016. doi: 10.1103/PhysRevE.93.052420.

[12] P. Venkatakrishnan, R. Sukanesh, and S. Sangeetha. Detection of quadratic phase coupling from human EEG signals using higher order statistics and spectra. Signal, Image and Video Processing, 5:217–229, 2011. doi: 10.1007/s11760-010-0156-x.

[13] Sarah Bartz, Christina Andreou, and Guido Nolte. Beyond pairwise interactions: The totally antisymmetric part of the bispectrum as coupling measure of at least three interacting sources. Frontiers in Neuroinformatics, 14:573750, 2020. doi: 10.3389/fninf.2020.573750.

[14] Laura Marzetti, Stefania Della Penna, Aaron Z. Snyder, Valentina Pizzella, Guido Nolte, Francesco de Pasquale, Gian Luca Romani, and Maurizio Corbetta. Frequency specific interactions of MEG resting-state activity within and across brain networks as revealed by the multivariate interaction measure. NeuroImage, 79: 172–183, 2013. doi: 10.1016/j.neuroimage.2013.04.062.

[15] Huixia He and David J. Thomson. Canonical bicoherence analysis of dynamic EEG data. Journal of Computational Neuroscience, 29(1-2):23–34, 2009. doi: 10.1007/s10827-009-0177-z.

[16] Alessandro E.P. Villa and Igor V. Tetko. Cross-frequency coupling in mesiotemporal EEG recordings of epileptic patients. Journal of Physiology-Paris, 104(3-4):197–202, 2010. doi: 10.1016/j.jphysparis.2009.11.024.

[17] Andreas K. Engel, Christian Gerloff, Claus C. Hilgetag, and Guido Nolte. Intrinsic coupling modes: Multiscale interactions in ongoing brain activity. Neuron, 80(4):867–886, 2013. doi: 10.1016/j.neuron.2013.09.038.

[18] Dana Lahat, Tülay Adali, and Christian Jutten. Multimodal data fusion: An overview of methods, challenges, and prospects. Proceedings of the IEEE, 103(9):1449–1477, 2015. doi: 10.1109/JPROC.2015.2460697.

[19] Surya Ganguli and Haim Sompolinsky. Compressed sensing, sparsity, and dimensionality in neuronal information processing and data analysis. Annual Review of Neuroscience, 35:485–508, 2012. doi: 10.1146/annurev-neuro-062111-150410.

[20] Carlos Oscar S. Sorzano, Javier Vargas, and Ana Pascual-Montano. A survey of dimensionality reduction techniques. arXiv preprint 1403.2877, 2014. URL https://arxiv.org/abs/1403.2877.

[21] John P. Cunningham and Zoubin Ghahramani. Linear dimensionality reduction: Survey, insights, and generalizations. Journal of Machine Learning Research, 16:2859–2900, 2015.

[22] John P. Cunningham and Byron M. Yu. Dimensionality reduction for large-scale neural recordings. Nature Neuroscience, 17:1500–1509, 2014. doi: 10.1038/nn.3776.

[23] Haidong Huang, Zhengming Ma, and Guokai Zhang. Dimensionality reduction of tensors based on manifold-regularized tucker decomposition and its iterative solution. International Journal of Machine Learning and Cybernetics, 13(2):509–522, 2021. doi: 10.1007/s13042-021-01422-5.

[24] Laurens J.P. van der Maaten and Geoffrey E. Hinton. Visualizing data using t-sne. Journal of Machine Learning Research, 9:2579–2605, 2008.

[25] Fengyu Cong, Qiu-Hua Lin, Li-Dan Kuang, Xiao-Feng Gong, Piia Astikainen, and Tapani Ristaniemi. Tensor decomposition of EEG signals: A brief review. Journal of Neuroscience Methods, 248:59–69, 2015. doi: 10.1016/j.jneumeth.2015.03.018.

[26] Dounia Mulders, Cyril de Bodt, Nicolas Lejeune, John A. Lee, André Mouraux and Michel Verleysen. Tensor factorization to extract patterns in multimodal EEG data. In Proceedings of the 27th European Symposium on Artificial Neural Networks, Computational Intelligence and Machine Learning (ESANN 2019), pages 601–606, Bruges, Belgium, 2019. i6doc.com. ISBN 978-287-587-065-0. URL https://www.esann.org/sites/default/files/proceedings/legacy/es2019-130.pdf.

[27] URL https://github.com/guidonolte/bsfit.

[28] URL https://www.uke.de/english/departments-institutes/institutes/neurophysiology-and-pathophysiology/research/research-groups/index.html.

[29] J. Douglas Carroll and Jih-Jie Chang. Analysis of individual differences in multidimensional scaling via an n-way generalization of ‘eckart-young’ decomposition. Psychometrika, 35(3):283–319, 1970. doi: 10.1007/BF02310791.

[30] Richard A. Harshman. Foundations of the parafac procedure: Models and conditions for an ‘explanatory’ multi-modal factor analysis. UCLA Working Papers in Phonetics, 16:1–84, 1970.

[31] Laura Marzetti, Cosimo Del Gratta, and Guido Nolte. Understanding brain connectivity from EEG data by identifying systems composed of interacting sources. NeuroImage, 42(1):87–98, 2008. doi: 10.1016/j.neuroimage.2008.04.250.

[32] Guido Nolte, Laura Marzetti, and Pedro Valdes-Sosa. Minimum overlap component analysis (MOCA) of EEG/MEG data for more than two sources. Journal of Neuroscience Methods, 183(1):72–76, 2009. doi: 10.1016/j.jneumeth.2009.07.006.

[33] Guido Nolte and George Dassios. Analytic expansion of the EEG lead field for realistic volume conductors. Physics in Medicine and Biology, 50(16):3807–3823, 2005. doi: 10.1088/0031-9155/50/16/010.

[34] Rakesh Ranjan, Bikash Chandra Sahana, and Ashish Kumar Bhandari. Ocular artifact elimination from electroencephalography signals: A systematic review. Biocybernetics and Biomedical Engineering, 41(3): 960–996, 2021. doi: 10.1016/j.bbe.2021.06.007.

[35] G. J. E. Duque, P. A. Munera, C. D. Trujillo, H. D. A. Urrego, and V. A. M. Hernandez. System for processing and simulation of brain signals. In 2009 IEEE Latin-American Conference on Communications (LATINCOM), pages 1–6, 2009. doi: 10.1109/LATINCOM.2009.5304853.

[36] Tonio Ball, Markus Kern, Isabella Mutschler, Ad Aertsen, and Andreas Schulze-Bonhage. Signal quality of simultaneously recorded invasive and non-invasive EEG. NeuroImage, 46(3):708–716, 2009. ISSN 1053-8119. doi: 10.1016/j.neuroimage.2009.02.028.

[37] Farzad Barzegaran, Thomas Kühnis, and Martin Meyer. EEGsourcesim: A framework for realistic simulation of EEG scalp data using mri-based forward models and dynamic source activation. Journal of Neuroscience Methods, 328:108395, 2019. doi: 10.1016/j.jneumeth.2019.108395.

[38] Giulia Cisotto. REPAC: Reliable Estimation of Phase-Amplitude Coupling in Brain Networks. In ICASSP 2021 - 2021 IEEE International Conference on Acoustics, Speech and Signal Processing (ICASSP), pages 1075–1079, Toronto, ON, Canada, 2021. IEEE. doi: 10.1109/ICASSP39728.2021.9414749.

[39] C. Andreou, G. Leicht, G. Nolte, N. Polomac, S. Moritz, A. Karow, I.L. Hanganu-Opatz, A.K. Engel, and C. Mulert. Resting-state theta-band connectivity and verbal memory in schizophrenia and in the high-risk state. Schizophrenia Research, 161(2-3):299–307, 2015.

[40] C. Andreou, G. Nolte, G. Leicht, N. Polomac, I.L. Hanganu-Opatz, M. Lambert, A.K. Engel, and C. Mulert. Increased resting-state gamma-band connectivity in first-episode schizophrenia. Schizophrenia Bulletin, 41(4):930–939, 2015.

[41] John C Mosher, Paul S Lewis, and Richard M Leahy. Multiple dipole modeling and localization from spatio-temporal meg data. IEEE transactions on biomedical engineering, 39(6):541–557, 1992.

[42] J.C. Mosher and R.M. Leahy. Recursive music: A framework for eeg and meg source localization. IEEE Transactions on Biomedical Engineering, 45(11):1342–1354, 1998. doi: 10.1109/10.725331.

[43] Forooz Shahbazi, Arne Ewald, and Guido Nolte. Self-consistent music: An approach to the localization of true brain interactions from eeg/meg data. NeuroImage, 112:299–309, 2015.

[44] Niko Mäkelä, Matti Stenroos, Jukka Sarvas, and Risto J Ilmoniemi. Truncated rap-music (trap-music) for meg and eeg source localization. NeuroImage, 167:73–83, 2018.

[45] Xiao-Liang Xu, Bobby Xu, and Bin He. An alternative subspace approach to eeg dipole source localization. Physics in Medicine & Biology, 49(2):327, 2004.

